# Modification of scale-free electrophysiological activity induced by changes in excitatory-inhibitory balance is governed by the dynamics of multiple oscillatory relaxation processes

**DOI:** 10.1101/203786

**Authors:** Suresh D Muthukumaraswamy, David TJ Liley

**Affiliations:** School of Pharmacy and Centre for Brain Research, Faculty of Medical and Health Sciences. The University of Auckland, Auckland, New Zealand; Centre for Human Psychopharmacology, School of Health Sciences, Swinburne University of Technology, Melbourne, Australia

**Author notes:** **Corresponding Author** Suresh Muthukumaraswamy, PhD, School of Pharmacy, Faculty of Medical and Health Sciences, Private Bag, 92019, Auckland, New Zealand.

**Keywords:** Magnetoencephalography,, Electroencephalography,, Electrocorticography,, Power Spectrum,, Scale-free activity

## Abstract

Neurophysiological recordings are dominated by arhythmical activity whose spectra can be characterised by power-law functions, and on this basis are often referred to as reflecting scale-free brain dynamics (1/*f* ^***β***^). Relatively little is known regarding the neural generators and temporal dynamics of this arhythmical behaviour compared to rhythmical behaviour. Here we used Irregularly Resampled AutoSpectral Analysis (IRASA) to quantify ***β***, in both the high (5-100 Hz, ***β***_hf_) and low frequency bands (0.1-2.5 Hz, ***β***_lf_ in EEG/MEG/ECoG recordings and to separate arhythmical from oscillatory modes of activity, such as, alpha rhythms. In MEG/EEG/ECoG data, we demonstrate that oscillatory alpha power dynamically correlates over time with ***β***_hf_ and similarly, participants with higher rhythmical alpha power have higher ***β***_hf_). In a series of MEG investigations using the GABA reuptake inhibitor tiagabine, the glutamatergic AMPA receptor antagonist perampanel, the NMDA receptor antagonist ketamine and the mixed partial serotonergic agonist LSD we reveal systematic effects of excitation-inhibition balance on both ***β***_hf_ and ***β***_lf_. Additionally, strong modulations of ***β***_hf_ are seen in monkey ECoG data during general anaesthesia using propofol and ketamine. Surrogate data analysis demonstrates that arhythmical activity is generated by both linear and non-linear schemes, with non-linear effects emerging at critical boundaries. We develop and test a unifying model which can explain, the 1/f nature of electrophysiological spectra, their dynamic interaction with oscillatory rhythms as well as the sensitivity of 1/f activity to excitation-inhibition balance by considering electrophysiological spectra as being generated by a collection of stochastically perturbed damped oscillators having a distribution of relaxation rates.

---

Numerous physical systems display scale-free (also termed fractal) temporal dynamics, characterised by power-law (1/*f* ^***β***^) spectral densities. Temporal signals displaying power-law behaviour have been observed throughout the nervous system at various spatial scales, from membrane potentials (El Boustani et al., 2009), single channel currents (Banerjee et al., 2006), and the spiking activity populations of neurons (Beggs and Plenz, 2003) through to the more global activities measured by electrocorticography, electroencephalography and magnetoencephalography (ECoG/EEG/MEG) (Dehghani et al., 2010; Freeman et al., 2003; Miller et al., 2009) and functional magnetic resonance imaging (He, 2011). The potential functional importance of power law scaling in the brain is underscored by its alteration in neuropsychiatric conditions (Lai et al., 2010; Maxim et al., 2005; Wei et al., 2013), ageing (Voytek et al., 2015; Wink et al., 2006), and its dynamic modification in task states (Barnes et al., 2009; Churchill et al., 2016; He, 2011; He et al., 2010; Oostenveld et al., 2011; Wink et al., 2006). Despite this, the physiologic mechanisms by which power-law scalings are generated is poorly understood and their significance remains controversial (Buzsaki, 2006; Buzsaki et al., 2012). In particular, it has been argued that the existence of power laws in the brain might indicate that the brain is in a state of self-organised criticality, where it is sensitively poised on a boundary between two qualitatively different dynamical states, potentially an optimal state for information transmission (Beggs and Timme, 2012; Beggs, 2008; Oostenveld et al., 2011). Alternatively, with respect to ECoG/EEG/MEG, it has been demonstrated that the 1/*f* properties of underlying local field potentials can potentially be explained by low-pass filtering of ionic current flow at dendritic processes, through the incorporation of their inherent geometric and capacitive properties (Linden et al., 2010; Pettersen and Einevoll, 2008). Similarly, it has been shown that 1/*f* power spectra can be created by ionic current flow in a convoluted extracellular space, if the media is considered as a complex arrangement of resistors and capacitors (Bedard and Destexhe, 2009; Bedard et al., 2006).

For decades, the dominant approach to the analysis of ECoG/EEG/MEG has been to consider either band-limited power changes, in the attempt to characterise brain rhythms, or evoked responses, while the background activity is often regarded as “1/*f* noise”. However, it has been noted that the 1/*f* activity forms the bulk of the content of the ECoG/EEG/MEG (Bullock et al., 1995; Freeman, 2004; He et al., 2010) and actually very little of the activity present in recordings with these modalities can mathematically be classified as rhythmical (Bullock et al., 2003). Indeed, even when oscillations do occur, they frequently have a non-sinusoidal morphology (Cole and Voytek, 2017). This is at odds with the view that the ECoG/EEG/MEG arises from the collective behaviour of a large set of independent oscillators, and instead is consistent with the idea of generators interacting in complex cross-frequency, often broadband patterns (Bullock et al., 1995). In our previous pharmaco-EEG/MEG studies we have observed that drug-induced changes of cortical power often show broadband spectral changes that cross the classically defined EEG frequency ranges, suggesting that underlying 1/*f* activity may be pharmacologically sensitive (Carhart-Harris et al., 2016; Kuhlmann et al., 2013; Muthukumaraswamy et al., 2015; Nutt et al., 2015).

Accurate estimation of the 1/*f* spectral component of the measurement of the ECoG/EEG/MEG, and in particular the value of ***β***, is however, non-trivial in empirical data. After transformation of the power spectrum to log-log coordinates ***β*** can in theory simply be estimated by standard least squares regression (P = 1/*f*^***β***^, ≡ log(P) = -***β***log(f)). However, as ECoG/EEG/MEG recordings are a compound mixture of non-stationary oscillatory rhythms, potentially with harmonic components, overlying the 1/*f* spectrum, these oscillations can substantially bias ***β*** estimation. Nevertheless, various investigators have provided estimates of ***β*** with the values obtained varying depending on the analysis technique used and the recording modality studied (Dehghani et al., 2010; He et al., 2010; Miller et al., 2009; Pritchard, 1992). In general, estimates of ***β*** lies in the range (0< ***β*** < 4), typically between 1 and 2 (between pink and brown noise), often with “knee” frequencies described, where the 1/*f* spectrum appears to deviate from linearity. Recently, Wen and Liu (Wen and Liu, 2016a, b) introduced a new method termed, Irregularly Resampled AutoSpectral Analysis (IRASA) which utilises non-integer/reciprocal resampling of a time-series to allow separate quantification of oscillatory and fractal components of the time-series and estimation of ***β***. In this work, we used IRASA to quantify the fractal component of the ECoG/EEG/MEG power spectra, focussing on ***β***, its interaction with oscillatory rhythms and its sensitivity to excitation-inhibition balance in the cerebral cortex. We demonstrate that the scalings observed and the changes subsequently induced in them pharmacologically can be economically accounted for if resting brain activity is viewed as arising from the collective behaviour of a distribution of alpha-band relaxation processes.

## Materials and Methods

### Description of data sources

Data were re-analysed from resting-state recordings from four pharmaco-MEG studies involving the drugs tiagabine, ketamine, perampanel and LSD (Carhart-Harris et al., 2016; Muthukumaraswamy et al., 2016; Muthukumaraswamy et al., 2015; Nutt et al., 2015). The reader is referred to those papers for comprehensive descriptions of the experiments and key features are to be found in Table S1. All whole-head MEG recordings were made using a CTF 275-channel radial gradiometer system sampled at 1200 Hz (0–300 Hz bandpass). An additional 29 reference channels were recorded for noise cancellation purposes and the primary sensors were analysed as synthetic third-order gradiometers (Vrba and Robinson, 2001).

Unpublished EEG data from 5 minute eyes-closed recordings were also analysed. The participants consisted of seventeen healthy male participants (mean age = 23). Participants were seated with their eyes-closed. 64 channel EEG was recorded continuously (1000 Hz sampling rate; 0.1 µV resolution, 250 Hz low pass filter) using Acticap Ag/AgCl active shielded electrodes with Brain Products MR Plus amplifiers (Brain Products GmbH, Munich, Germany). All electrode impedances were typically maintained below 10 kΩ. EEG was acquired using an FCz reference and a ground electrode located at AFz.

Monkey electrocorticographic data were downloaded from the publicly available neurotycho database (http://neurotycho.org/) (Nagasaka et al., 2011). From these data, we selected recordings from two monkeys, “George” and “Chibi”. Both these monkeys had data from two sessions of propofol anaesthesia (~5 mg/kg intravenous propofol) and another two sessions with ketamine anaesthesia (~5.1 mg/kg intramuscular) available for analysis. There were approximately 15 minutes of eye-open and closed data and 9 minutes of anaesthesia data for analysis.

### MEG pre-processing

Pre-processing of data was performed using custom scripts based on the Fieldtrip toolbox (Oostenveld et al., 2011). Continuous MEG data were first visually inspected and those sections of data with gross artefacts were removed from the data. Datasets were excluded if data were not of acceptable quality (e.g. excessive artefact sections). Independent component analysis (ICA), as implemented in Fieldtrip/ EEGLAB was then performed on the remaining data sections. Visual inspection was used to identify components related to ocular, cardiac and muscular artefacts and these components were then projected out of the data. Source modelling of the data was performed using the Fieldtrip toolbox (Oostenveld et al., 2011). For each participant individual forward models were generated from their individual structural MRI scan using a “single-shell” model (Nolte, 2003). Due to computational complexity of subsequent processing with IRASA, in order to reduce the data, an atlas-based beamformer approach was used (Hillebrand et al., 2012). Broadband virtual sensor time-series were constructed using a linearly constrained minimum variance beamformer with 5% regularisation (Van Veen et al., 1997) at 90 cortical and subcortical seed locations as specified in the automated anatomical labelling atlas (Tzourio-Mazoyer et al., 2002). Frequency spectra were calculated and inspected for each dataset to “sanity-check” the source solutions.

### EEG pre-processing

Similar to the MEG data, continuous EEG data were first visually inspected and those sections of data with gross artefacts were removed. Datasets were excluded if data were not of acceptable quality (e.g. excessive artefact sections). Bad channels were also removed at this stage and then ICA used to remove artefacts. As individual MRI scans were not available no source modelling on these data were performed. After artefact removal, data were average-referenced and any missing channels were replaced using spherical-spline interpolation (Perrin et al., 1989).

### ECoG pre-processing

Three sections of each recording day were considered for analysis, resting eyes-open, eyes-closed and anaesthesia. These sections were extracted from the data using annotated event-markers. Each dataset contained 128 electrodes and using the provided electrode maps, 64-channel bipolar montages were computed to reduce any far-field artefact contamination (Fukushima et al., 2014; Yanagawa et al., 2013).

### IRASA (Irregularly Resampled AutoSpectral Analysis) computation

The IRASA technique developed by Wen et al (Wen and Liu, 2016b) and applied in (Wen and Liu, 2016a) allows for the separation in the frequency domain of the fractal and oscillatory components of a time-series. In this sense it is similar to coarse-grained spectral analysis (CGSA) (Yamamoto Y and RL, 1993), although IRASA is superior in situations, where harmonics may exist in the data, or where complex interactions between the oscillatory and fractal components may be present. Such interactions may well be the case with neural time-series (Wen and Liu, 2016b) as our data will demonstrate. The reader is referred to (Wen and Liu, 2016b) for a full mathematical treatment of IRASA and a MATLAB implementation can be found online (https://engineering.purdue.edu/libi/lab/Resource.html). Nevertheless, a brief description of the algorithm is provided here. In the IRASA algorithm, each epoch of data to be analysed is divided into 15 overlapping and evenly distributed sub-epochs that each comprise 90% of the total length of the original epoch. For each sub-epoch of data, the power spectrum is calculated using a Fast Fourier Transform (FFT). Prior to FFT computation, data is tapered with a Hanning window and a length of FFT is used such that it is the next power of two after multiplying the sub-epoch data length by the largest resampling value to be used (*h*(max) -see later). The average spectral power of the sub-epochs is taken as an estimate of the total (mixed) power spectrum. The sub-epochs are then resampled using a range of resampling factors (*h*) and their reciprocals (1/*h)*. We set *h*=1.1 to 2.9 in steps of 0.05 with *h*≠2. The sub-epochs are upsampled for one of the reciprocal pairs, using cubic spline interpolation and are downsampled with a similar interpolation scheme, but following low-pass filtering to avoid aliasing for the other reciprocal pair. The mean of each upsampled sub-epoch for a given *h,*1/*h* pair is calculated, and then the median across *h* values is calculated as a representation of the fractal power for the epoch. The oscillatory power spectrum is then calculated as the difference between the total and fractal power spectrum. The IRASA process is depicted in Figure S1. It should be noted that the primary free parameter is the choice of resampling factors *h*. A larger range of *h* provides better suppression of residual oscillatory power in the fractal power spectrum, at the expense of a reduction in the highest frequency available for analysis. Given that all the datasets we analysed were sampled at 1000Hz or greater, with an *h*(max)=2.9, this left a maximum resolvable frequency of 250 Hz. Unless otherwise specified, data were divided into sections of 10s length ready prior to computation of IRASA. The length of the data epoch sets the lowest frequency available for analysis. For epochs of 10s duration sampled at 1,000Hz this is 0.03 Hz. We note that while Liu and Wen (Wen and Liu, 2016a, b) set *h* =1.1 to 1.9 in steps of 0.05 we found in some of our data cases, those settings led to residual spectral mixing (identifiable as a bump in the log-log spectra). Hence a broader range for *h* was used here. In order to estimate the power law exponent (b), the fractal power spectrum was transformed to log-log coordinates and linear regression performed. To avoid biasing regression estimates towards the higher frequencies, where more sampling points exist in logarithmic space, frequency estimates are resampled to be evenly spaced in logarithmic coordinates prior to computation of the regression.

### Statistical analyses

All linear mixed-effects modelling was performed in MATLAB using the fitlme.m function and initially verified with SPSS®. Participants were considered as random effects and time as a fixed effect. Throughout, statistical comparison of models was performed against null distributions obtained from 5000 resamplings. The individual plotted locations from the AAL Atlas corresponded to: Cuneus = Cuneus_L (Index = 45), Precentral = Precental_L (Index = 1), Frontal = Frontal_Sup_Medial_L (Index = 23), Temporal = Temporal_Mid_L (Index = 85).

### Autoregressive-moving-average (ARMA) modelling of alpha-band (8-13 Hz) activity

In order to obtain smoother estimates of single-trial alpha-band activity parametric ARMA modelling was used. Here we down-sampled data to 80 Hz and fitted an (8,3) ARMA model to the data using the ARMASA MATLAB toolbox (Broersen, 2002), according to:

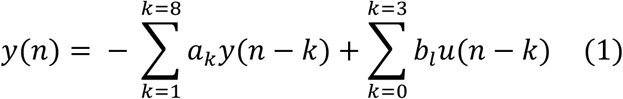

where y(n) is the observed signal at time n and *a_k_* and b_k_ are the (time-varying) autoregressive (AR) and moving-average (MA) parameters. u(n) represents a stationary white-noise process. The AR and MA orders used are derived from a physiologically plausible mean field model of the mammalian spontaneous electroencephalogram (Liley et al., 2010) and accord well with empirical determinations of optimal autoregressive (range, 3 – 14) and moving average (range, 2 – 5) orders obtained from resting awake eyes-closed EEG using a range of information theoretic criteria (Liley et al., 2010). From the AR and MA parameters autocorrelation functions (and hence the power spectral density) can be estimated. Further, following z transformation:

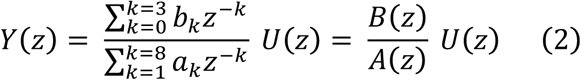

where solutions to A(z)= 0 will give the system poles and solutions to B(z)=0 give the system zeros. The poles correspond to the dominant oscillations in the signal or the peaks in the power spectrum of the signal, while the zeros essentially indicate frequencies where power is nulled. Together this allowed several parameters to be extracted for alpha (8-13Hz) including, peak frequency, power at peak frequency (peak power), the integrated power in the 8-13Hz band (band power) and full-width at half-maximum (FWHM). A parameter estimating alpha-band “damping” (−*σ*_*l*_) was estimated directly from the AR parameters. The damping parameter quantifies how quickly the oscillation will decay over time after it has been perturbed and the boundary parameter where no power is present. Specifically, *σ*_*l*_ was calculated as *f_s_* ln|*z*|, where *f_s_* is the sampling frequency and |*z*| the average modulus of all poles having a frequency between 8 and 13Hz

## Results

MEG data were typically analysed in the source space (Figure 1a), which not only permits more powerful inferences about source locations but helps to suppress both low and high-frequency artefacts (Muthukumaraswamy, 2013). IRASA was then used to estimate ***β*** from the fractal part of the power spectrum (Figure 1b). Similar to others, we observed knee frequencies in the data and hence divided the fits into a low frequency component denoted ***β***_lf_ (0.1-2.5 Hz) and a high frequency component ***β***_hf_ (5-100 Hz unless otherwise specified). Figure 1c) shows comparable data and fits can be obtained in MEG sensors and includes an example empty room noise recording. Our MEG sensors generally showed a white noise profile (>1Hz) in contrast to some MEG system sensors which show more of a pink profile to nearly 100 Hz (see (Hamalainen et al., 1993) Figure 30). Below 0.1 Hz signals were around the noise floor so this was used as a lower-bound analysis frequency.

**Figure 1:**
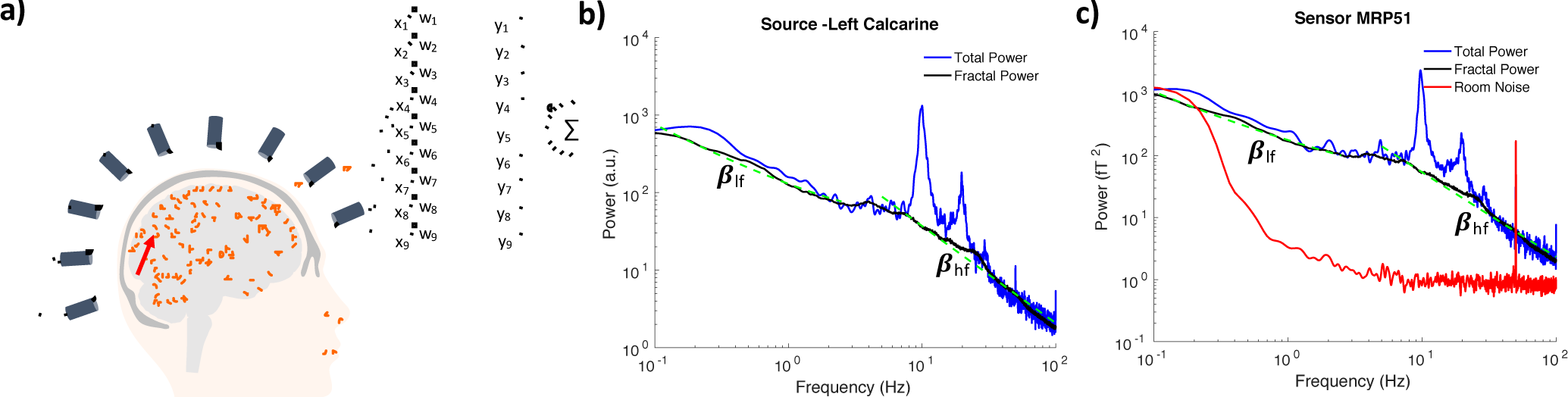
Illustration of the IRASA process for MEG data analysis. a) A single example source dipole (red arrow) in the left calcarine cortex is recorded from multiple MEG detectors (only 9/273 shown for illustrative purposes). A Linearly Constrained Minimum Variance (LCMV) beamformer is illustrated, whereby sources can be reconstructed as a linear weighting of all the recorded signals. Data can be analysed in either the source space following beamforming (b) or as recorded at sensors c). Total power is plotted in blue and fractal power in black. Estimates of **β**_hf_ and **β**_lf_ are obtained by fitting a straight line (dashed green lines) to the fractal power spectrum obtained following IRASA. For the sensor space data an empty room noise recording is plotted in red in c) and shows a white noise spectrum (**β** =~0) in contrast to the brain where **β**>0.5.

### Correlation between rhythmical and non-rhythmical activity

***β***_hf_ and ***β***_*lf*_ values obtained from resting eyes-closed data showed considerable variation across the cortex. For the high frequencies (***β***_hf_) mean slope was 1.21 (range 0.78-1.45) while for the lower frequencies the mean slope was 0.76 (range = 0.56-0.98). A clear spatial pattern was evident (Figures 2a and 2b), such that in the higher frequencies steeper slopes are present in posterior areas whereas for the lower frequencies (***β***_lf_) steeper slopes are present in the frontal cortex. Given the spatial overlap of the higher frequencies with alpha generating areas, oscillatory alpha power (8-13 Hz) was extracted for each 10s epoch and correlated with ***β***_hf_. This analysis (Figure 2 c and d) reveal the existence of striking temporal correlations, indicating a dynamic relationship between alpha power and the underlying fractal power in the higher frequency band. Here we computed ***β***_hf_ over the 20-100 Hz range. This result was replicated in data from three further studies (Figures S2, S3, S4) including both eyes-open and eyes-closed data. To confirm the broad applicability of the correlation between and alpha power and ***β***_hf_, its existence was confirmed first in EEG data (Figure S5) and then in monkey ECoG data (Figure 3 - see Figure S6 for underlying ECoG spectra). For eyes-closed ECoG data the mean ***β***_hf_ was 1.96 (range 1.33-2.76) whereas for eyes-open data, the mean ***β***_hf_ was 1.85 (range 1.30-2.76). For eyes-closed data the mean ***β***_lf_ was 0.57 (range -0.53-2.13) whereas for eyes-open data, mean ***β***_lf_ was 0.64 (range -0.22-2.05). It was noted that ***β***_hf_ values were generally highest and ***β***_lf_ lowest in occipital areas, implying less overall linearity in the scaling of the spectrum in these areas. A variety of pre-processing approaches were used for the different recording modalities, suggesting that the observed correlations are highly robust. We conducted multiple numerical simulations to confirm this effect was not an artefact of the IRASA analysis procedure itself by producing artificial 1/*f* data with embedded alpha oscillations at a range of signal-to-noise levels, ***β***_hf_ values and variances (see Figures S7, S8).

**Figure 2:**
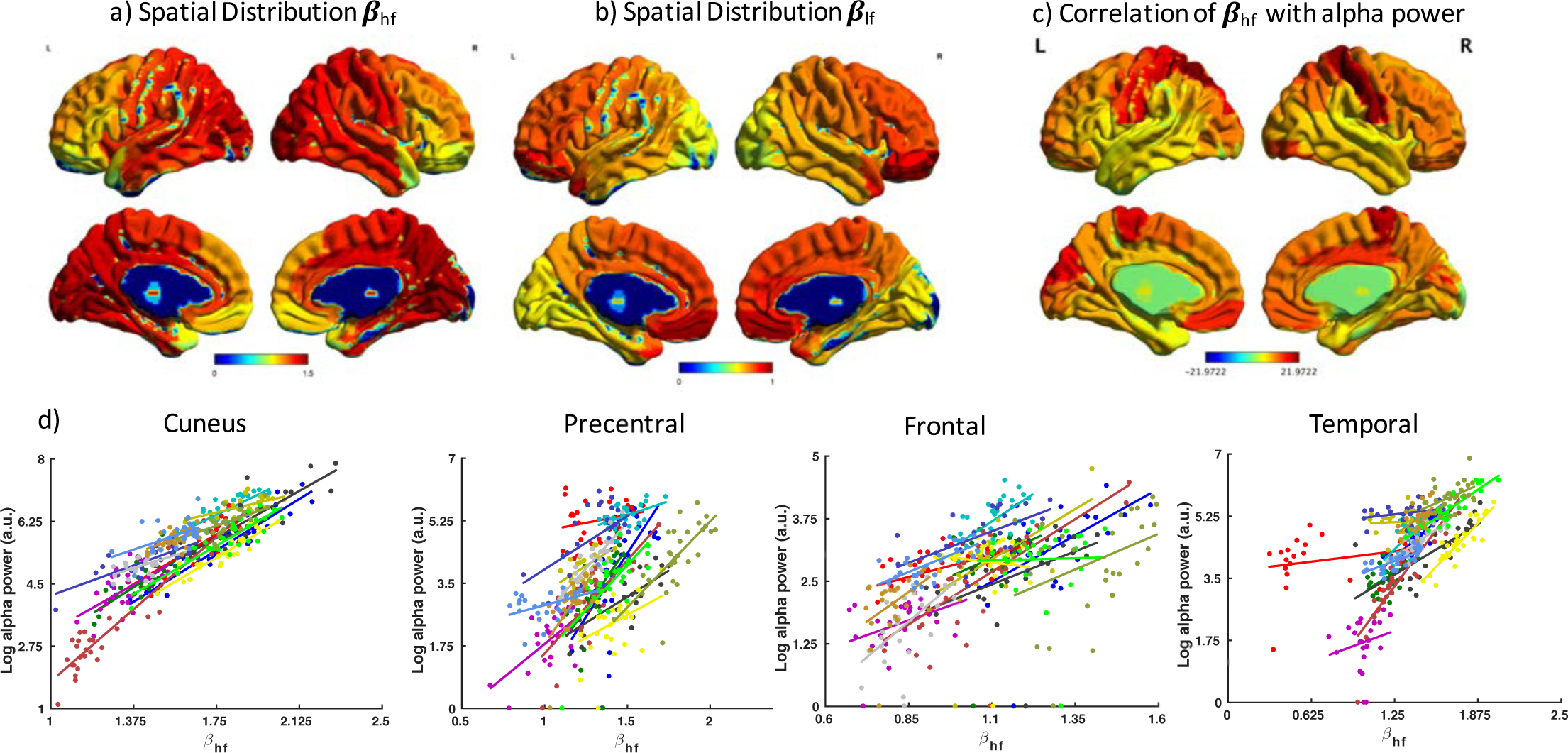
a and b) Estimates of **β**_hf_ and **β**_lf_ averaged across participants for each of the 90 cortical locations of the AAL (automated anatomical labelling) atlas (Tzourio-Mazoyer et al., 2002) for the eyes-closed resting placebo from the tiagabine experiment. Here **β**_hf_ was estimated from 20-100 Hz and **β**_lf_ from 0.1-2.5Hz. c) Temporal correlations of **β**_hf_ with oscillatory alpha power. Plotted are the t statistic values from a linear mixed model fitting all data, which was permuted 5000 times to obtain null distributions. The false discovery rate was then used to correct the images for multiple comparisons (p<.05). d) Data from a selection of left cortical areas including: cuneus, precentral gyrus, frontal cortex, and temporal cortex are plotted. Each participant (n=15) is plotted with a different colour with individual data points and regression lines included.

**Figure 3:**
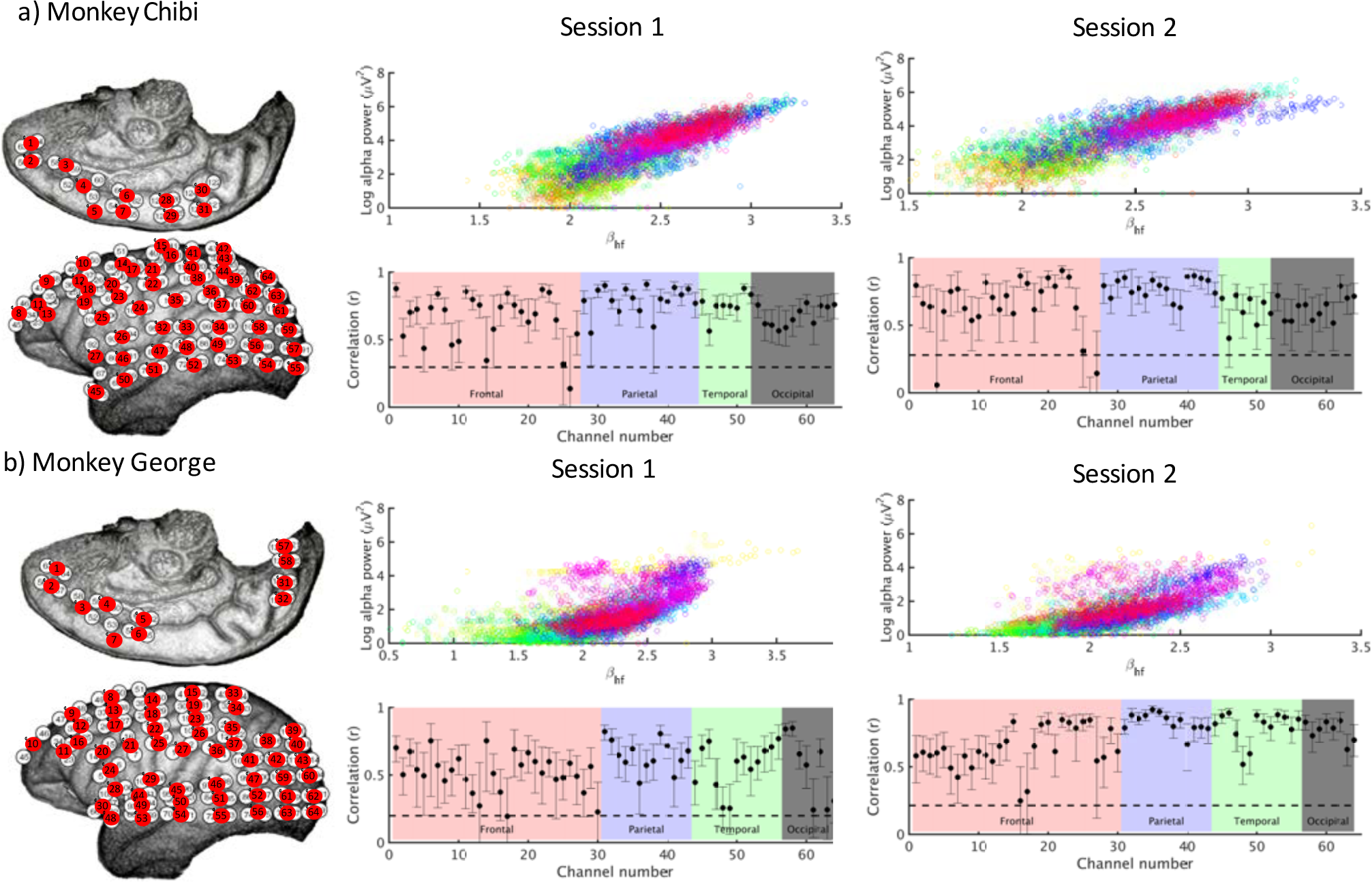
Temporal correlations of **β**_hf_ (20-100 Hz) with oscillatory alpha power (8-13 Hz for ECoG data from two monkeys, Chibi a) and George b). Data analysed were from two resting eyes-closed recordings made on separate days. The left hand panels show electrode positions for each monkey, who each had 128 electrodes implanted in the left hemisphere (white circles). These were analysed as 64 bipolar electrode pairs (red circles). The upper scatter plots display estimates of alpha power versus **β**_hf_ for each 10s epoch of data with each bipolar electrode plotted as a separate colour. The lower scatter plots show the Pearson correlation coefficient (r) for each electrode with bootstrapped confidence intervals (5000 repeats) for each correlation provided. The dashed line represents the FDR corrected significance threshold for that session. Electrodes have been broadly allocated to each of the four lobes.

Given the temporal correlation that exists within participants we examined whether across participants a similar correlation exists. Inter-participant correlations were found such that those participants who exhibit high alpha power have higher ***β***_hf_ values (Figures S9, S10, S11, S12, S13). Given the dynamical temporal relationships between ***β***_hf_ and alpha power, we investigated, but did not find, evidence of lag-lead effects using 1s data windows stepped forward in 100 ms bins (Figure S14). Correlations between ***β***_lf_ and alpha power were explored, but we found little evidence for such a relationship (Figure S15).

### Excitation-inhibition balance determines non-rhythmical activity

The potential for ***β***_hf_ and ***β***_*lf*_ values to be modified by pharmacological intervention was investigated in four pharmaco-MEG studies involving the GABA reuptake inhibitor tiagabine, the non-selective NMDA receptor antagonist ketamine, the selective non-competitive AMPA antagonist perampanel and the neuromodulator and serotonergic hallucinogen LSD. All drugs were given at relatively low doses, which mostly caused sedation, apart from LSD which caused hallucinations (Carhart-Harris et al., 2016). Significant modification of ***β***_hf_ and ***β***_lf_ values were seen for tiagabine, perampanel, ketamine and LSD (Figure 4). For tiagabine, concurrent increases of ***β***_hf_ (placebo mean=1.21, range=0.78-1.45; tiagabine mean=1.66, range=1.29-1.91) and decreases in ***β***_lf_ values (placebo mean =0.76, range 0.56-0.99; tiagabine mean = 0.43, range 0.09- 0.72) are evident across the cortex. With tiagabine, large increases in oscillatory rhythms in the delta, theta and lower alpha bands can also be seen. For perampanel, modest local increases of ***β***_hf_ (placebo mean=1.22, range=0.87-1.46; perampanel mean=1.25, range=0.88-1.51) and local decreases in ***β***_lf_ values (placebo mean =0.72, range 0.61-0.84; perampanel mean = 0.67, range 0.59-0.85) can be seen. With perampanel, increases in oscillatory delta, and alpha band activity can be seen with a slight slowing of peak alpha frequency. Conversely, for ketamine, decreases in ***β***_hf_ (placebo mean=1.14, range=0.83-1.35; ketamine mean=1.03, range=0.75-1.23) and increases in ***β***_lf_ values (placebo mean=0.77, range=0.66-0.94; ketamine mean=0.87 range= 0.78- 0.99) are evident. Although not significant across the cortex, this trend appeared in every brain area but did not always pass correction for multiple comparisons. Reductions in delta and alpha oscillatory activities can also be seen with ketamine. In terms of ***β***_hf_ and ***β***_lf_, similar to ketamine, LSD showed decreases in ***β***_hf_ (placebo mean=1.25, range=0.94-1.45; LSD mean=0.92, range=0.51-1.10) and increases in ***β***_lf_ values (placebo mean=0.74, range=0.66-0.94; LSD mean=0.95, range=0.86-1.05) across the cortex. In the oscillatory domain, a decrease in power, but acceleration of alpha oscillatory frequency can be seen (Walter, 1957).

**Figure 4:**
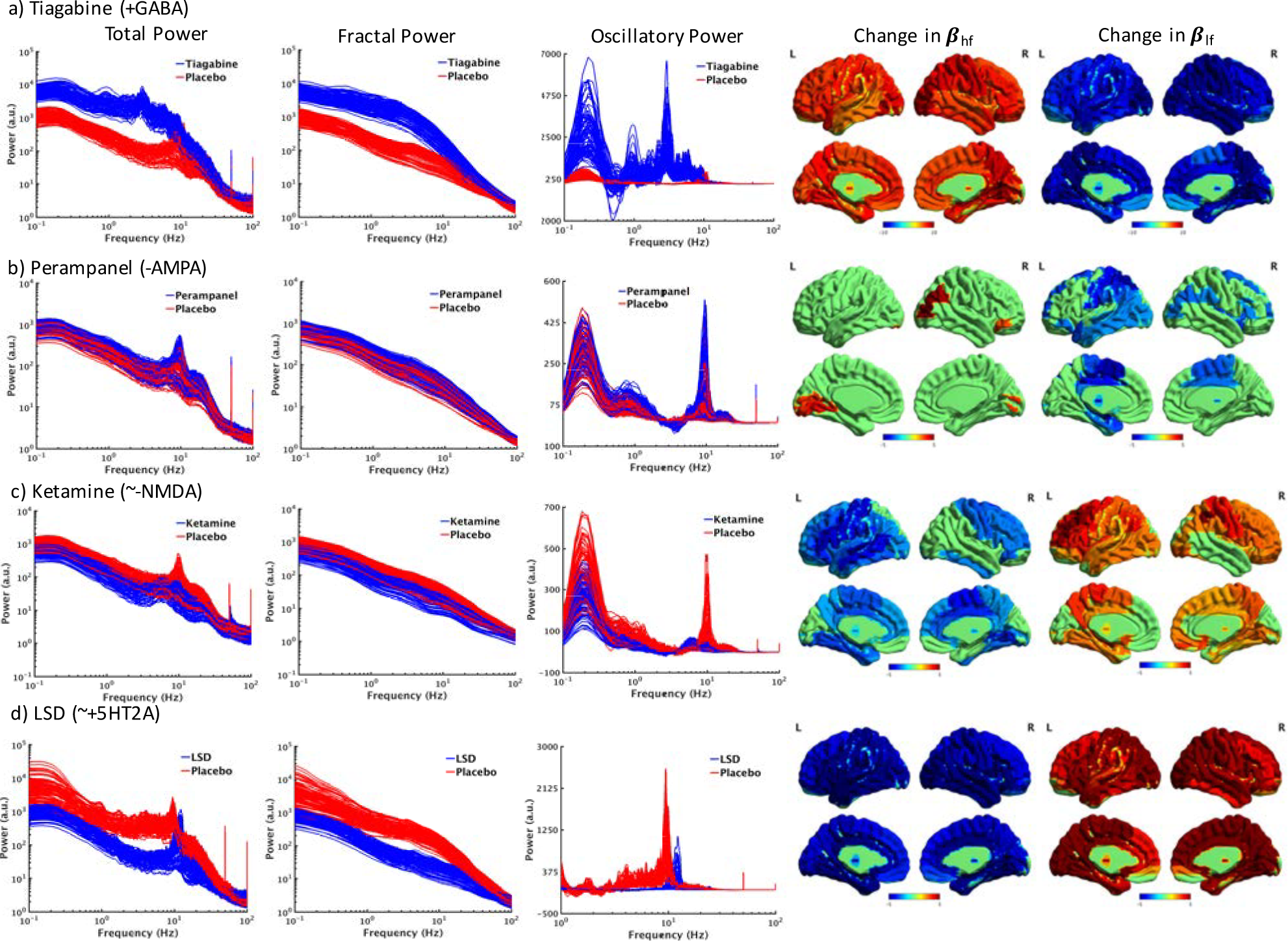
IRASA analysis of resting pharmaco-MEG data from four studies using a) tiagabine, b) perampanel, c) ketamine and d) LSD. The left hand sets of spectra show the total power spectrum from 90 cortical locations of the AAL atlas for drug and placebo in log-log coordinates, while the centre spectra show just the fractal part of the power spectra in log-log coordinates. The right hand spectra show the residual oscillatory activity with only the x axis in log space. Note the LSD axis was truncated at 1Hz to allow better visualisation of the alpha peak frequency shift. The brain plots show statistical comparison of fitted **β**_hf_ (5-100 Hz) and **β**_lf_ (0.1-2.5 Hz) fitted to the fractal power spectrum. Red colours indicate an increase in **β** and blue colours a decrease, between drug and placebo condition. Data shown are from t tests compared with a null distribution via permutation testing (5000 permutations) at each location, corrected for multiple comparisons using the false discovery rate (p<.05). Note: a) is scaled to t=±10 whereas b), c) and d) are scaled to t=±5.

Traditionally, quantitative EEG analysis, particularly in pharmaco-EEG is used to describe the absolute and relative power in each of pre-defined frequency bands. However, quantification of spectral power with IRASA allows subdivision into both fractal and oscillatory activity. For each of the MEG studies, a typical quantitative MEG/EEG analysis was performed (see Figures S16, S17 S18, S19). We note that the division of the spectrum into fractal and oscillatory modes can reveal new insights into pharmacologically-induced spectral patterns, which are more physiologically interpretable than relative EEG power.

Given the effects seen in pharmaco-MEG data, we investigated whether similar effects exist in monkey ECoG data – here during anaesthesia induced by propofol and ketamine (Figure 5). Strong changes in ***β*** estimates were seen during these sessions, however, there was significantly more regional variation than with the pharmaco-MEG data. The most consistent effects were in the propofol ***β***_hf_ parameter estimates which showed anterior increases and posterior decreases. Clearly, averaging ***β*** estimates across brain regions was not appropriate for these data. Considerable differences were also seen between the two monkeys. However, the results across 15 the two sessions for each monkey were highly consistent for propofol (***β***_hf_ for Chibi R2=0.98; ***β***_hf_ for George R2=0.97; ***β***_lf_ for Chibi R2=0.73; ***β***_lf_ for George R2=0.60) and relatively so for ketamine (***β***_hf_ for Chibi R2=0.95; ***β***_hf_ for George R2=0.62; ***β***_lf_ for Chibi R2=0.45; ***β***_lf_ for George R2=0.39).

**Figure 5:**
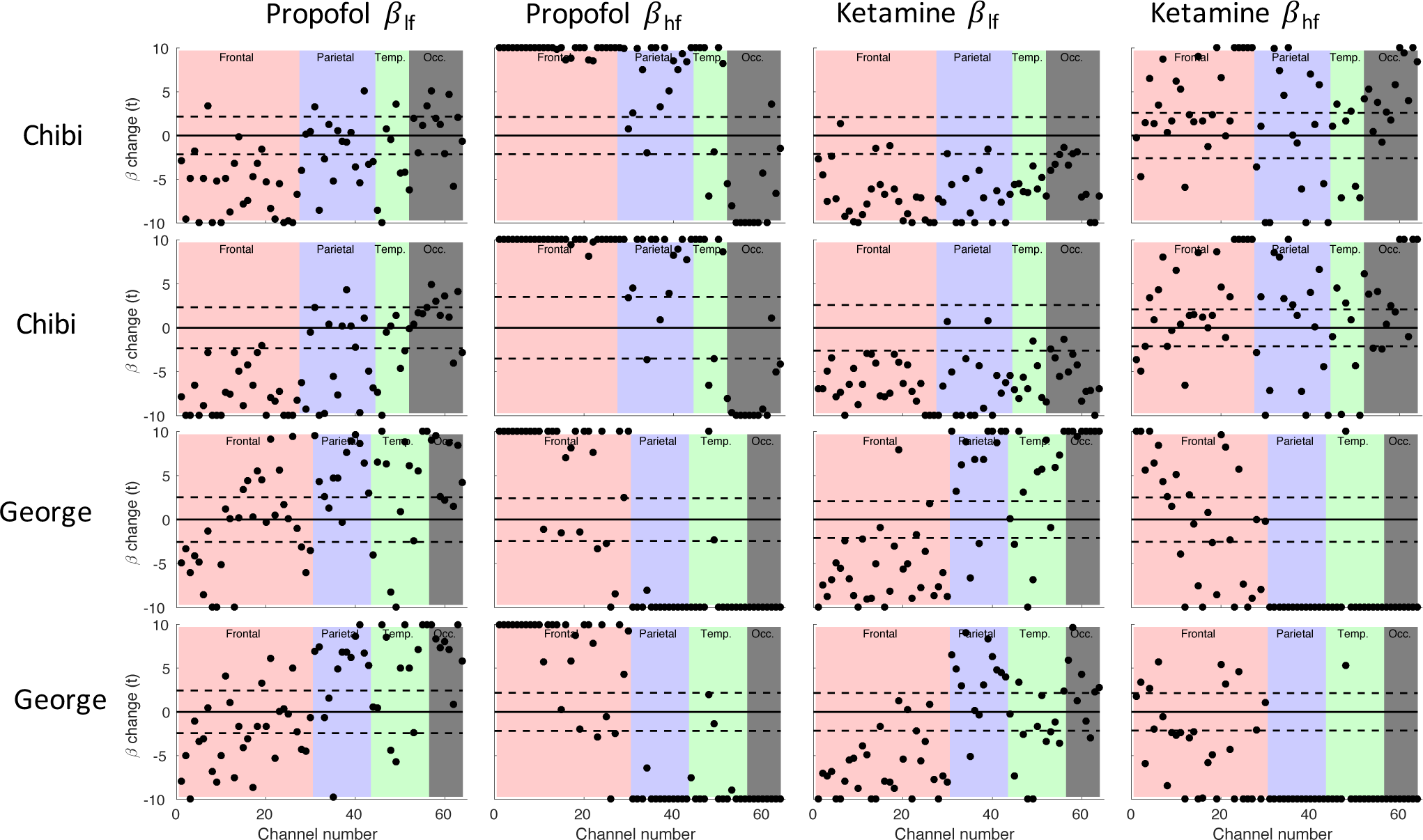
IRASA analysis of ECoG data for a total of four sessions from two monkeys (George and Chibi). The four sessions consisted of two sessions with propofol and two sessions with ketamine. Each column provides estimates of **β**_hf_ or **β**_lf_. Estimates are z scores for estimates of the change in **β** from anaesthesia compared to preceding eyes-closed data. For convenience of visualisation where t>10, t=10 or where t<-10, t=-10 in plots. The dashed line indicates the FDR threshold for the z score in each session. Individual p value estimates were obtained by comparison with a null distribution obtained from 5000 resamplings. Background colours provide the cortical areas with channel maps found in Figure 3.

### Autoregressive modelling of excitation-inhibition balance modifications

Initial attempts to correlate ***β***_hf_ with alpha peak frequency and spectral width showed no relationships. However, this may have been caused by noisy parameter estimation due to using nonparametric (Fourier) spectral estimation. More accurate estimates of alpha full width at half maximum (FWHM) and peak frequency were made using fixed-order autoregressive moving average (ARMA) modelling (Figure 6 and S20). These showed that alpha-frequency FWHM is strongly inversely correlated to alpha power and hence also to ***β***_hf_. The FWHM for alpha activity from the ARMA analysis was typically around 2 Hz. Further, because the FWHM of a spectral resonance in a white-noise driven damped linear oscillatory system is known to be proportional to the temporal damping of the resonance, we also calculated the damping (−σ_l_, see middle panel Figure 7 and Methods) of the alpha band oscillatory modes (poles) identified by our fixed-order ARMA analysis. We found that alpha band power, total band power and ***β***_hf_ were inversely correlated with damping (Figure 6a-b). We further investigated the relationships between ***β***_hf_, alpha damping, and alpha power in each of the four pharmaco-MEG intervention experiments (Figure 6c-f). In general, it can be seen in both drug and placebo states ***β***_hf_ is correlated with alpha power, and alpha power is dependent on the degree of damping. A variety of drug-induced parameter estimations can be seen. Tiagabine (Figure 6c) increases ***β***_hf_ and increases damping, whereas LSD (Figure 6d) decreases ***β***_hf_ with little change in damping. Conversely, ketamine (Figure 6e) decreased ***β***_hf_ but increased damping while perampanel (Figure 6f) caused a small increase in ***β***_hf_ with a small decrease in damping. See Table S3 for more details.

**Figure 6:**
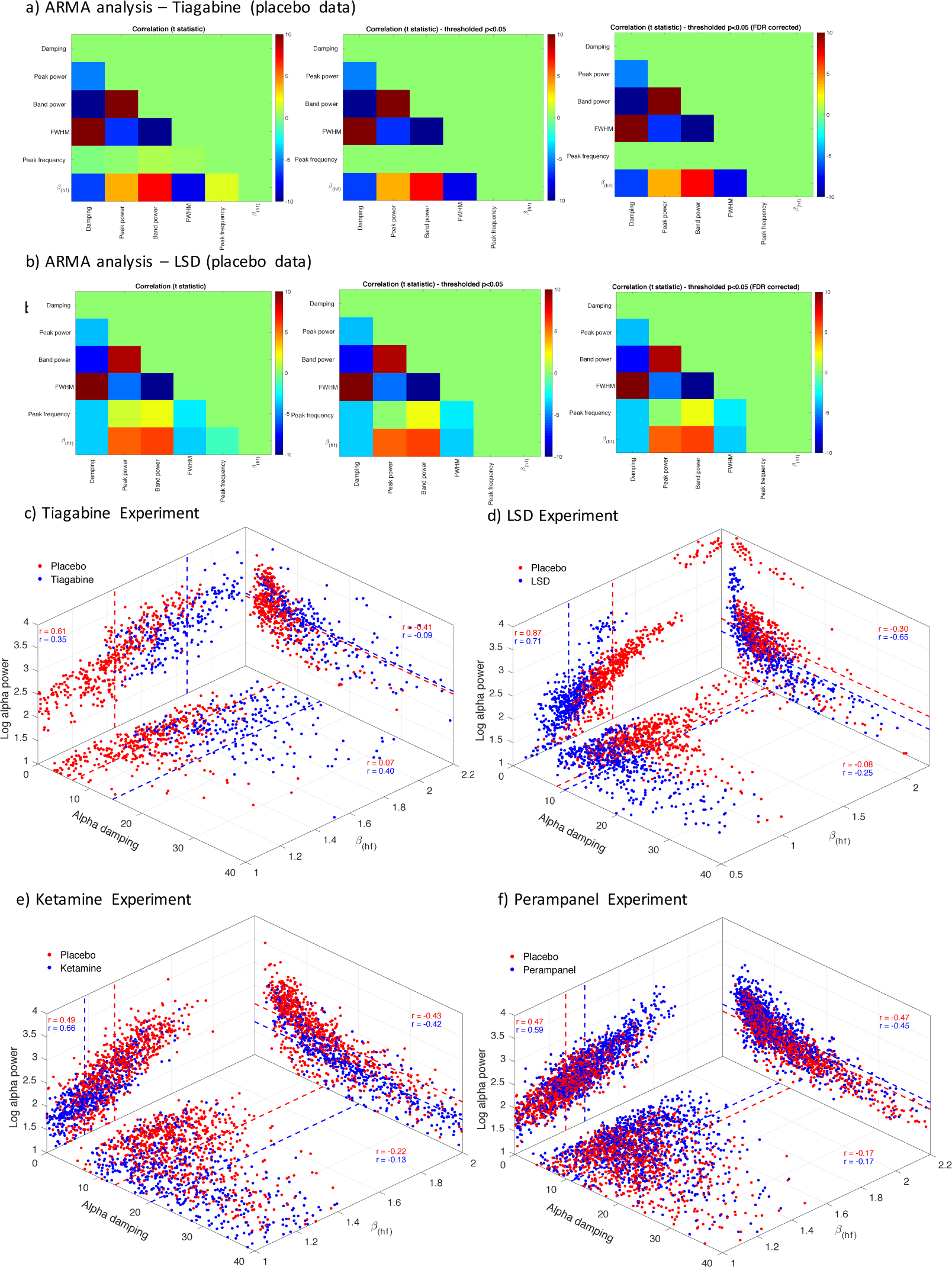
Results of ARMA analysis (see also Figure S16) for all participants for the tiagabine (a) and LSD (b) experiments. A single brain location rich in oscillatory alpha (left cuneus) was selected for analysis. From the ARMA analysis, a number of parameters were quantified for the alpha-band (8-13 Hz), including peak frequency, power at peak frequency (peak power), the integrated power in the 8-13 Hz band (band power) and FWHM. A parameter estimating alpha-band damping was estimated directly from the AR parameters. The damping parameter quantifies how quickly an oscillation will decay over time after it has been perturbed. Correlations were performed for each of these parameters and also **β**_hf_ using a mixed-model regression, recomputed 5000 times to obtain a null distribution for comparison. The results demonstrate that **β**_hf_ is positively correlated to metrics of alpha power and inversely correlated to FWHM and damping. As one might predict, damping and FWHM are inversely correlated to power metrics. c-f). The relationship between alpha power (here computed by IRASA/FFT), alpha damping and **β**_hf_ using posterior channels from Figure 4 that showed large pharmacologically induced **β**_hf_ changes. Fixed effect statistics are presented with mean parameter estimates indicated with dashed lines. Statistical comparisons across conditions can be found in Table S2.

### A mechanistic model of 1/f electrophysiological activity based on damped oscillators

Given the observed relationships between ***β***_hf_, alpha pole damping and power this suggests a mechanistic explanation of electrophysiological spectra based on a collection of relaxation processes. Because it is often empirically difficult to establish that spontaneous ECoG/EEG/MEG results from an identifiable non-linear process it is often sufficient to describe its dynamical behaviour in terms of a noise driven linear system having a number of characteristic oscillatory modes. One of the simplest such possible characterisations involves assuming the existence of two damped oscillations – one having a frequency close to zero and another having a frequency of alpha (f_γ_). In this case the resulting power spectrum is equal to the sum of two Lorentzians, each of which is parameterised by the damping and frequency of the respective oscillatory modes. This simple model can be further generalised to the case in which there are multiple alpha modes having a distribution of dampings. We note from (Milotti, 2002) that for a relaxation process having a frequency *f*α and a distribution of dampings (*p*(*σ*)) the spectral density is given by:

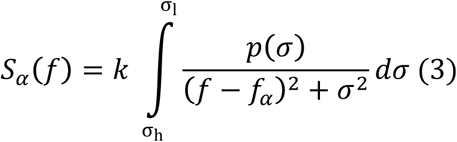

where *k* >0 is a constant that takes into account the recording arrangement and the electrical properties of the skull, scalp and meninges. To enable exact integration for the subsequent analytical calculation of a derivative with respect to frequency (see Model fitting of electrophysiological spectra in Methods) we assume a trapezoidal distribution for *p*(*σ*):

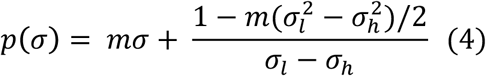

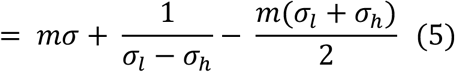

where |*m*| ≤ 2(*σ_h_* − *σ_l_*)^−2^ is the slope over the range [σ_h_, σ_l_]. From Equations 3-5 it can be seen that the EEG spectral shape can be controlled by four parameters, alpha peak frequency *f*_α_, the damping probability distribution limits (σ_l_ and σ_h_) and m (when m=0 the distribution reverts to a uniform distribution). In real data, ARMA analysis allows both σ_l_ and *f_α_* to be explicitly measured where σ_l_ is the most weakly damped pole (see Figure 7a). *m* and σ_h_ are obtained by constrained fitting. In order to fit our model, Equation 3 was integrated to reveal an estimate of the power spectrum:

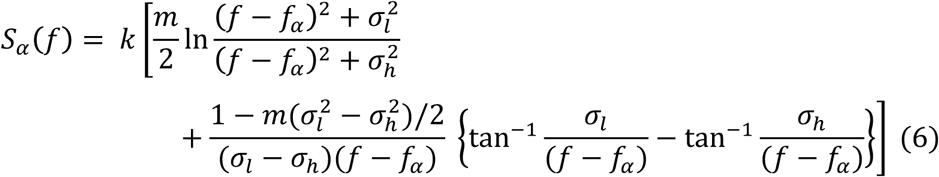

To estimate *m* and σ_h_ we define the log-log frequency scaling of *S_α_*(*f*), 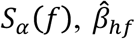, as the slope of log_10_ *S*_α_(10^*F*^) evaluated at the logarithm of the midpoint frequency over which β_*hf*_ was empirically estimated, i.e.

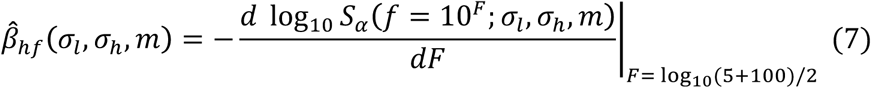

where the dependence on *σ_l_*, *σ_h_*, *m* is made explicit. On the basis that the most weakly damped alpha modes, *σ_l_*, can be estimated directly from the ARMA analysis and under the simplest assumption that the distribution of dampings are uniform (i.e. *m* = 0) solutions to 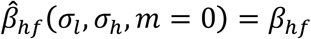 will enable us to uniquely estimate *σ_h_* (see Table S3). In the case that no solution can be found for *m* = 0 a non-uniform distribution of dampings was assumed. However, this results in an under-determined problem and thus additional constraints were applied in order to find a solution. The constraint we choose to apply is one that minimises the Kolmogorov-Smirnov distance between a prior uniform placebo distribution, as estimates of *σ_h_* can be found for all placebo conditions for *m* = 0, and a resulting posterior distribution for a drug condition that arises as a solution to 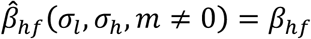.

**Figure 7a.**
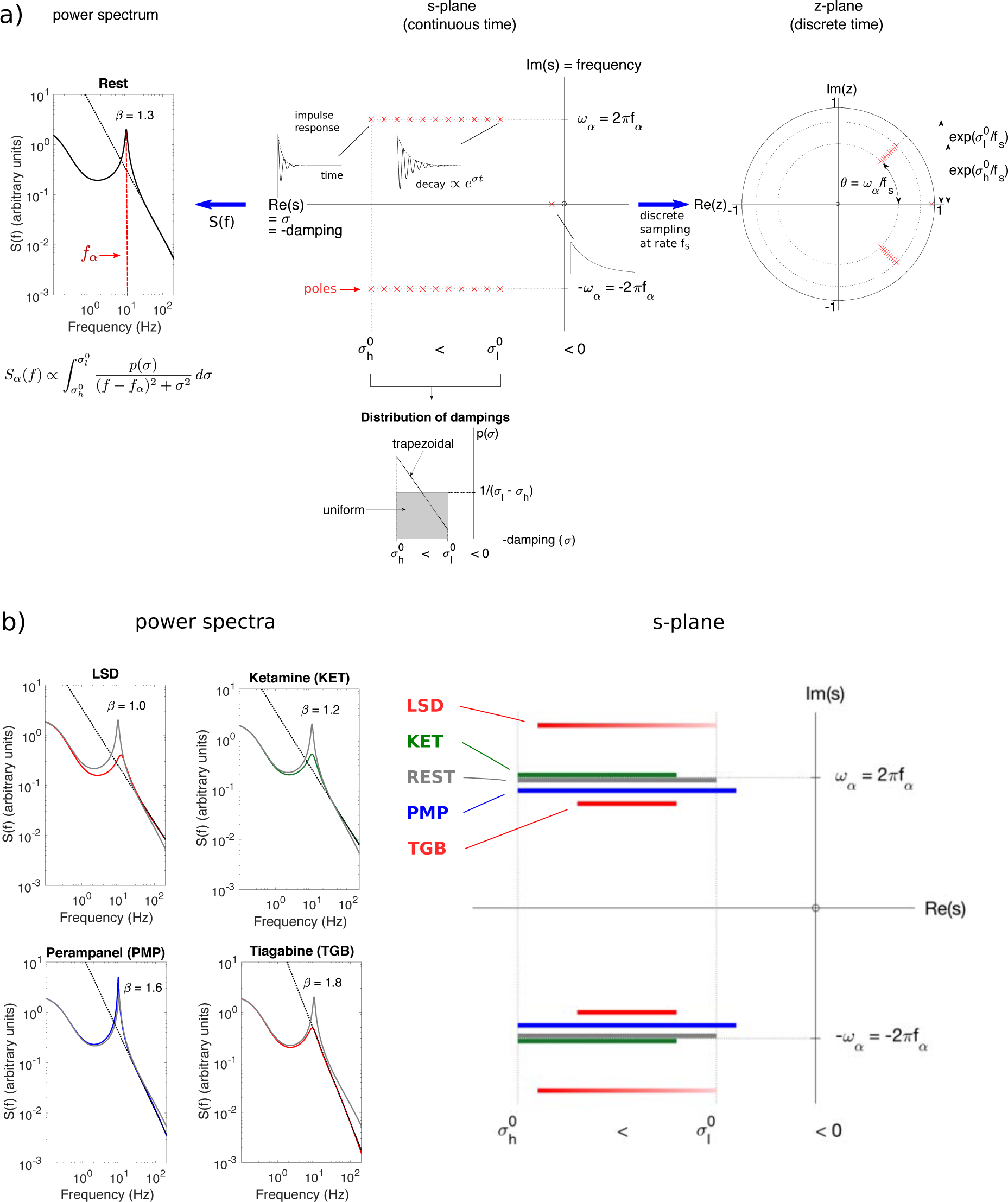
A model of electrophysiological spectra. In the middle figure a set of individual oscillatory modes are specified in terms of their co-ordinates in the complex s-plane (red crosses), where the impulse responses associated with a number of these oscillatory modes have also been shown. More generally these oscillatory modes are called poles, and in terms of the underlying linear system correspond to the eigenvalues of one or more linear evolution matrices. Nonoscillatory modes are represented by a single pole whereas oscillatory modes require a pole and its complex conjugate. By assuming that this linear system so specified is moment to moment randomly perturbed, the resulting power spectrum will be the sum of the frequency responses associated with each of the poles. This is illustrated in the left-hand figure for a model resting spectrum. The spectral peaks at f = 0 and f = f_α_ correspond to the values of the two imaginary co-ordinates for the respective poles in the middle figure, whereas the power scaling for f > f_α_ reflects the distribution of dampings (lower subfigure of the middle figure) associated with the poles of frequency f_α_. Parametric time series methods can be used to empirically characterise the linear system illustrated in the middle figure and are able to provide additional information regarding dynamical structure not easily attained using non-parametric Fourier Transform methods. Auto Regressive-Moving Average (ARMA) time-series models are able to provide economical characterisations of weakly stochastic systems such as the MEG/EEG and allow estimation of σ_l_ and f_α_. Because AMRA analysis depends on the discrete sampling, at a rate f_s_, of a continuous time series the resulting parameterisation of the identified oscillatory modes is in terms of co-ordinates in the z-plane (right-hand figure). In this case frequency corresponds to the polar angle of the complex pole and damping to the negative of the natural logarithm of its radius. b) On the basis of empirically calculated changes in β_hf_ and alpha band damping (−σ_l_) various pharmacological agents are concluded to differentially affect the distribution of decay rates associated with alpha band relaxation processes. Perampanel (solid blue line) action is associated with the augmentation of the most-weakly damped alpha band relaxation processes. In contrast both ketamine (dashed blue line) and tiagabine (dashed red line) attenuate these weakly damped processes, but have opposite effects on heavily damped modes. LSD (solid red line) modifies the distribution slope (m) and accordingly is associated with an increase in the proportion of heavily damped alpha modes. The location of the bars on the imaginary axis reflects the empirically measured peak frequency of alpha (f_α_) relative to the resting spectrum.

From Equation (6) we note that

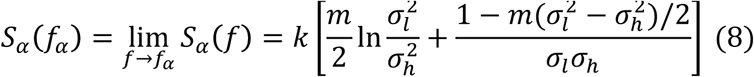

which for *m* = 0 reduces to *S_α_* (*f_α_*) = *k*/(*σ_l_ σ_h_*), thus allowing the possibility of also estimating *σ_h_* from empirical estimates of *S_α_* (*f_α_*) for known values of *k*, given that *σ_l_* can be estimated from the ARMA analysis of Equation (4). However, because *k* is in general not known we prefer to note that 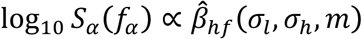 (see Figure S24) thus accounting for the empirical relationships identified between alpha power and *α_hf_* of Figure 6. Based on these equations we are able to account for the mechanistic generation of (1/f) electrophysiological spectral data for both eyes open and closed data as well as during pharmacological manipulation of the spectra by tiagabine, ketamine, LSD and perampanel (see Methods, Figure 7b and Table S3).

### Nonlinearity Testing

To test whether the generation of ***β***_hf_ relies on underlying nonlinear mechanisms we subjected a subset of the recorded data to nonlinearity testing using surrogate data generated using the iterated amplitude fast Fourier transform method (IAFFT). These analyses (Figure 8, Table S4) revealed that approximately 39% (1060/2740) of the placebo ***β***_hf_ estimates tested could be attributed to non-linearity at *α* = 0.05, with these estimates tending to fall at the more extreme values of ***β***_hf_. Nonlinearity was not observed in ***β***_hf_ estimates from synthetically produced data (pink noise with embedded oscillations (see Figure S21 and S22 for an outline of the method used). Across drugs we found that LSD significantly decreased the number of epochs that were nonlinear whereas perampanel increased the number of epochs that were nonlinear, while ketamine and tiagabine had no effect (see Table S4).

**Figure 8:**
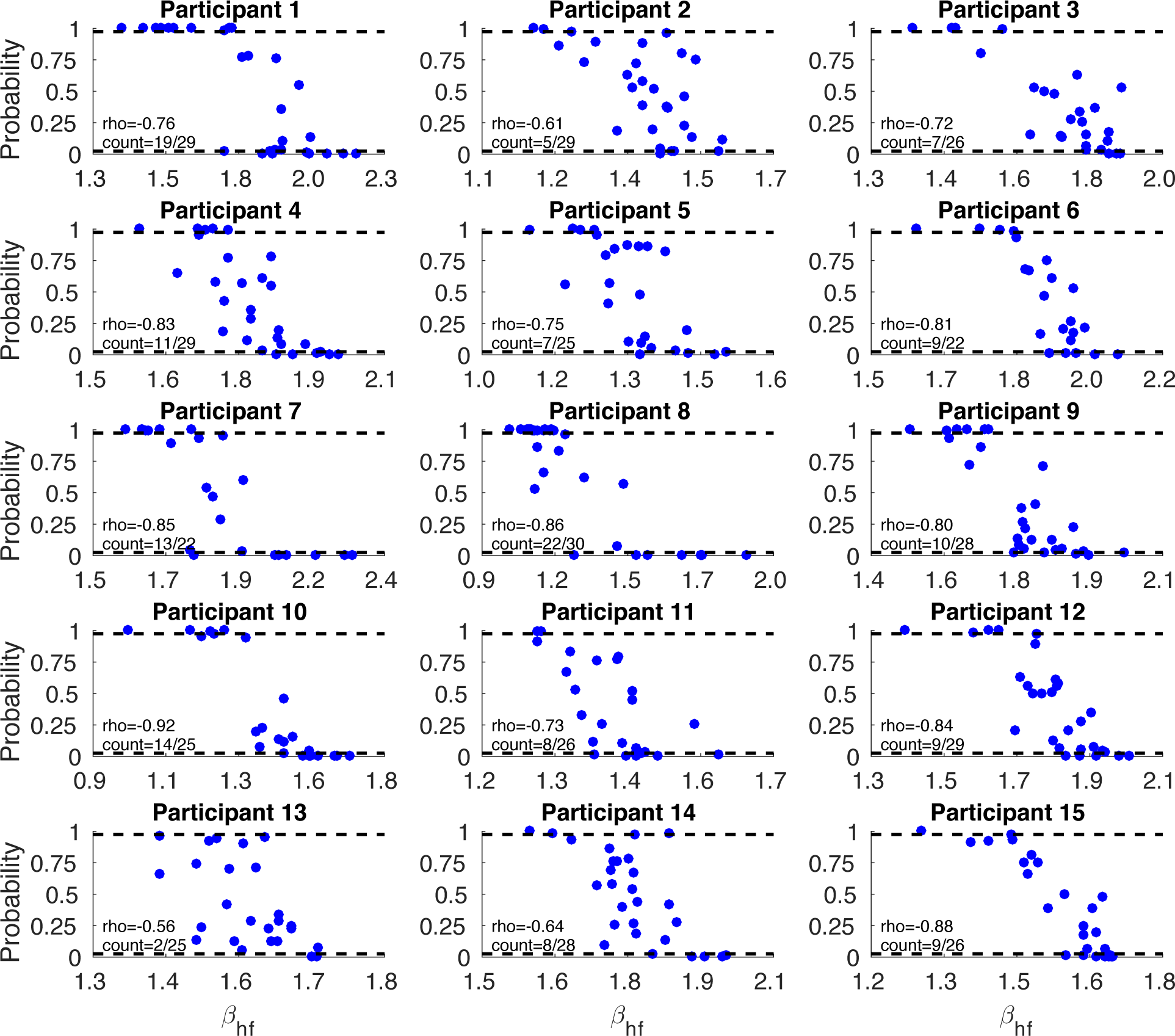
Results of surrogate data (nonlinearity) testing for a subset of the data (fifteen participants from placebo recordings from the tiagabine experiment – left cuneus. 1000 surrogate versions of each epoch were generated for each epoch using the IAFFT method (Schreiber and Schmitz, 1996) and **β**_hf_ calculated for both the original data and the surrogates. Location of obtained estimates are represented as a probability on the surrogate distribution. Those obtained estimates falling outside the 95 % (both tails) distribution are considered nonlinear. Counts of nonlinearity are provided for each participant with the overall count being 153/399 (38%) of epochs. We observed a clear effect where the nonlinear epochs appeared to lie more at the extreme values of **β**_hf_. To quantify this effect, Spearman’s rho is provided on each plot.

## Discussion

In these analyses we revealed for the first time striking dynamic correlations between oscillatory alpha power and scale-free ***β***_hf_ activity in ECoG/EEG/MEG in both humans and monkeys. Contrary to prevailing views (He, 2014) we find that scale-free arrhythmic brain electromagnetic activity can be robustly explained without the need to invoke mechanistically uncertain processes such as self-organized criticality and fractal temporal dynamics. Analysis using theoretically motivated, but empirically plausible, fixed-order ARMA models revealed that ***β***_hf_ was significantly inversely correlated with the damping of linear oscillatory modes in the alpha band during resting conditions (Figure 4). The existence of this inverse correlation suggests a clear explanation for the relationship between alpha power and ***β***_hf_. In the simplest situation, consider a simple damped harmonic oscillator in which the damping, and thus the FWHM and power, of the single oscillatory mode can be modified. Reductions in the damping of this system will lead to higher amplitude oscillations in the time domain and a narrowing of the spectral resonance in the frequency domain. The narrowing of this spectral resonance will be associated with a steeper fall off in power for higher frequencies. Therefore, the observed changes in alpha power can potentially be understood as the result of alterations in the damping of a single oscillatory mode having a resonant frequency within the alpha band. In our model, by a simple expansion of this to a more physiologically realistic probability distribution of damped oscillatory poles, a rich array of spectral behaviour can be explained. On this basis we are able to formally derive a relationship (see Methods) between ***β***_hf_ and the damping of alpha oscillatory activity and show that they are negatively correlated (Figure 6). The model we propose is sufficient to explain normal resting data as well as data from four interventional studies which showed various combinations of oscillatory alpha power and ***β***_hf_ changes. It may be that further expansion of this model is required when other rhythms are included such as (coupled) beta/theta rhythms. This could be accommodated by allowing more oscillatory modes and/or more complex damping probability distributions. However, for the data under consideration the current model was sufficient to explain all the data while allowing exact solutions.

Our interpretation is supported by the results of a number of relatively recent mean field modelling approaches aimed at understanding the rhythmogenesis of the resting mammalian electroencephalogram and in particular the alpha rhythm (Liley et al., 2002; Robinson et al., 2002). Such models are formulated in terms of the feedforward and feedback interactions of cortical excitatory and inhibitory neuronal populations and are parameterised in such a way that the effects of alterations in the strength and time course of synaptic connectivity on the emergent dynamical activity can be systematically investigated. While these models are typically constituted in terms of coupled non-linear partial differential equations, under certain conditions they admit near linear solutions. For example, the model of Liley et al. (Liley et al., 2002) suggests that the resting mammalian EEG can explained by the superposition of two noise driven damped linear resonances – one near 0 Hz and the other in the alpha band – with the damping of these resonances being determined by the strength and characteristic time course of synaptic excitation and inhibition. Our clear empirical observation of a knee frequency between ***β***_lf_ and ***β***_hf_, as well as the ability drugs affecting neurotransmission to modify ***β***_hf_, is consistent with such an interpretation. For small values of the damping, a state of near-criticality is predicted such that there is the possibility that small changes in input (external or internal) to the cortical neuronal populations may push the system over a critical boundary such that spatiotemporally organised non-linear activity emerges. The identified non-linearity of data epochs exhibiting extremal values of ***β***_hf_ (Figure 8) deviating from the mean ***β***_hf_ is broadly consistent with such a supposition. Nevertheless, we have not explicitly considered the role that non-linear processes play in the generation of the observed power law spectral behaviour. Further, we assumed that the relaxation processes arise solely from the decay of a perturbed linear system. We have not investigated the possibility that non-linear relaxation processes, such as the decay of a limit cycle oscillation within its basin of attraction, might also contribute to our empirical estimates and their subsequent pharmacological modification, as such decay would also be expected to contribute to the spectral broadening of the fundamental limit cycle oscillation frequency. Clearly such a situation, which falls outside of the scope of this current communication, needs to be studied.

Previously it has been shown that estimates of ***β*** are increased in older adults (Voytek et al., 2015) and decreased in patients with schizophrenia (Peterson et al., 2017). Hitherto, such differences between population groups could potentially be explained away by potential physiological differences in tissue filtering properties of the extracellular media, which can modify ***β*** estimates (Bedard and Destexhe, 2009; Bedard et al., 2006). However, tissue filtering alone cannot explain our results where we see modifications that occur within participants, both dynamically and following pharmacological interventions. As such, our work suggests an alternate interpretation for the modification of ***β*** estimates seen in special population groups. That is, ***β*** differences may reflect modifications of cortical excitation-inhibition balance on alpha pole damping. For ageing, most likely this could be attributed to decline in the GABA system with age. Several studies have noted reduced GABA concentration in humans (Porges et al., 2017), and monkey (He et al., 2016) measured with MRS, as well as reduced oscillatory gamma-frequency in humans (Porges et al., 2017), which is thought to depend on GABAergic mechanisms. Functionally-related decreased proportions of GABA interneurons have been observed in cat visual cortex (Hua et al., 2008) and indeed GABA agonists can restore visual function in ageing monkeys (Leventhal et al., 2003).

In these results we found generalised decreases in ***β***_hf_ and increases of ***β***_lf_ with ketamine in humans, whereas while the ECoG data in monkeys showed strong effects on these parameters, these were more variable in terms of spatial location, direction and across animal. Notably, the human data was collected at subanaesthetic doses (0.3 mg/kg) whereas the anaesthesia data is obtained at doses ~5 mg/kg. While ketamine is primarily considered an NMDA antagonist, it has a mixed pharmacology at clinically relevant concentrations, including but not limited to, actively inhibiting hyperpolarisation-activated cyclic nucleotide (HCN) channels, facilitating delta and mu opioid activity as well increasing release of dopamine and noradrenaline (see (Sleigh et al., 2014)). These mixed activities become increasingly relevant at increasing concentrations, for example, the EC50 of ketamine for NMDA channels is 1-5 *μ*M and slightly higher for HCN channels (8-16 *μ*M) (Chen et al., 2009). HCN channels, which mediate a depolarising inward current in response to hyperpolarisation, might be particularly relevant, given that HCN knockout mice show twofold less sensitivity to the hypnotic actions of ketamine (Chen et al., 2009). Similarly, propofol also causes inhibition (Chen et al., 2009) of HCN with decreased propofol sensitivity observed in HCN knockout mice. Moreover, neural field modelling of the EEG suggests that much of the effects of ketamine and propofol on EEG rhythmicity can be accounted for by their activity at HCN channels rather than primary NMDA/GABAergic effects (Bojak et al., 2013). By contrast, perampanel and tiagabine have much more selective pharmacological effects on excitation and inhibition respectively.

Some of our analyses to date, like much of the extant literature, have largely considered alpha rhythms as a somewhat stationary process, however, clearly this is not the case. In electrophysiological recordings, alpha rhythms emerge as bursts of varying amplitude, duration at irregular intervals. Given the clear relationships between alpha power, damping and scale-free activity, we are currently pursuing a more detailed examination of these dynamics. Similarly, since transient evoked responses exist in similar low-frequency bands it would be worth investigating their covariation (if any) with scale-free activity. As mentioned, previous work has demonstrated that scale-free, 1/f activity can be modified by task states (He, 2011; He et al., 2010) and can be altered in special populations suggesting functional relevance (Peterson et al., 2017; Voytek et al., 2015). Our data, demonstrate clear dynamical interactions between broadband scalefree activity and the more frequently studied oscillatory phenomena - interactions which are at odds with idea that electrophysiological activity is generated by many independent oscillators with different frequencies. Further, our explanatory model suggests that 1/f generation can be more simply explained by considering electrophysiological activity as a stochastic distribution of non-stationary damped oscillators. As such, the power law nature of macroscopic-scale electrophysiological spectra can be explained by relatively well understood physical phenomena and more complex explanations such as self-organised criticality need not be invoked. Given the macroscopic scale of our electrophysiological measurements we are unable to conclude whether individual relaxation oscillations are generated by circumscribed populations of cortical neurons. In the case that they are not, the recorded collective behaviour may instead reflect the superposition of correlated neuronal activity in multiple co-existing networks in a manner consistent with fluctuations in a population of Hebbian cell assemblies.

## Acknowledgements

Suresh Muthukumaraswamy is supported by a Rutherford Discovery Fellowship administered by the Royal Society of New Zealand. The authors thank Levin Kuhlmann for assistance with the ARMA modelling.

## References

Banerjee, J., Verma, M.K., Manna, S., Ghosh, S., 2006. Self-organised criticality and 1/f noise in single-channel current of voltage-dependent anion channel. Europhysics Letters 73, 457–463.

Barnes, A., Bullmore, E.T., Suckling, J., 2009. Endogenous human brain dynamics recover slowly following cognitive effort. PLoS One 4, e6626.

Bedard, C., Destexhe, A., 2009. Macroscopic models of local field potentials and the apparent 1/f noise in brain activity. Biophys J 96, 2589–2603.

Bedard, C., Kroger, H., Destexhe, A., 2006. Does the 1/f frequency scaling of brain signals reflect self-organized critical states? Phys Rev Lett 97, 118102.

Beggs, J., Timme, N., 2012. Being Critical of Criticality in the Brain. Frontiers in Physiology 3.

Beggs, J.M., 2008. The criticality hypothesis: how local cortical networks might optimize information processing. Philosophical Transactions of the Royal Society A: Mathematical, Physical and Engineering Sciences 366, 329–343.

Beggs, J.M., Plenz, D., 2003. Neuronal avalanches in neocortical circuits. J Neurosci 23, 11167–11177.

Bojak, I., Day, H.C., Liley, D.T., 2013. Ketamine, Propofol, and the EEG: A Neural Field Analysis of HCN1-Mediated Interactions. Front Comput Neurosci 7, 22.

Broersen, P.M.T., 2002. Automatic Spectral Analysis With Time Series Models. IEEE TRANSACTIONS ON INSTRUMENTATION AND MEASUREMENT 51, 211–216.

Bullock, T.H., McClune, M.C., Achimowicz, J.Z., Iragui-Madoz, V.J., Duckrow, R.B., Spencer, S.S., 1995. Temporal fluctuations in coherence of brain waves. Proc Natl Acad Sci U S A 92, 11568–11572.

Bullock, T.H., McClune, M.C., Enright, J.T., 2003. Are the electroencephalograms mainly rhythmic? Assessment of periodicity in wide-band time series. Neuroscience 121, 233–252.

Buzsaki, G., 2006. Rhythms of the Brain. Oxford University Press.

Buzsaki, G., Anastassiou, C.A., Koch, C., 2012. The origin of extracellular fields and currents—EEG, ECoG, LFP and spikes. Nat Rev Neurosci 13, 407–420.

Carhart-Harris, R.L., Muthukumaraswamy, S., Roseman, L., Kaelen, M., Droog, W., Murphy, K., Tagliazucchi, E., Schenberg, E.E., Nest, T., Orban, C., Leech, R., Williams, L.T., Williams, T.M., Bolstridge, M., Sessa, B., McGonigle, J., Sereno, M.I., Nichols, D., Hellyer, P.J., Hobden, P., Evans, J., Singh, K.D., Wise, R.G., Curran, H.V., Feilding, A., Nutt, D.J., 2016. Neural correlates of the LSD experience revealed by multimodal neuroimaging. Proceedings of the National Academy of Sciences USA 113, 4853–4858.

Chen, X., Shu, S., Bayliss, D.A., 2009. HCN1 channel subunits are a molecular substrate for hypnotic actions of ketamine. J Neurosci 29, 600–609.

Churchill, N.W., Spring, R., Grady, C., Cimprich, B., Askren, M.K., Reuter-Lorenz, P.A., Jung, M.S., Peltier, S., Strother, S.C., Berman, M.G., 2016. The suppression of scale-free fMRI brain dynamics across three different sources of effort: aging, task novelty and task difficulty. Sci Rep 6, 30895.

Cole, S.R., Voytek, B., 2017. Brain Oscillations and the Importance of Waveform Shape. Trends Cogn Sci 21, 137–149.

Dehghani, N., Bedard, C., Cash, S.S., Halgren, E., Destexhe, A., 2010. Comparative power spectral analysis of simultaneous elecroencephalographic and magnetoencephalographic recordings in humans suggests non-resistive extracellular media. J Comput Neurosci 29, 405–421.

El Boustani, S., Marre, O., Behuret, S., Baudot, P., Yger, P., Bal, T., Destexhe, A., Fregnac, Y., 2009. Network-state modulation of power-law frequency-scaling in visual cortical neurons. PLoS Comput Biol 5, e1000519.

Freeman, W.J., 2004. Origin, structure, and role of background EEG activity. Part 2. Analytic phase. Clin Neurophysiol 115, 2089–2107.

Freeman, W.J., Holmes, M.D., Burke, B.C., Vanhatalo, S., 2003. Spatial spectra of scalp EEG and EMG from awake humans. Clin Neurophysiol 114, 1053–1068.

Fukushima, M., Saunders, R.C., Mullarkey, M., Doyle, A.M., Mishkin, M., Fujii, N., 2014. An electrocorticographic electrode array for simultaneous recording from medial, lateral, and intrasulcal surface of the cortex in macaque monkeys. J Neurosci Methods 233, 155–165.

Hamalainen, M., Hari, R., Ilmoniemi, R.J., Knuutila, J., Lounasmaa, O.V., 1993. Magnetoencephalography - Theory, Instrumentation, and Applications to Noninvasive Studies of the Working Human Brain. Reviews of Modern Physics 65, 413–497.

He, B.J., 2011. Scale-free properties of the functional magnetic resonance imaging signal during rest and task. J Neurosci 31, 13786–13795.

He, B.J., 2014. Scale-free brain activity: past, present, and future. Trends Cogn Sci 18, 480–487.

He, B.J., Zempel, J.M., Snyder, A.Z., Raichle, M.E., 2010. The temporal structures and functional significance of scale-free brain activity. Neuron 66, 353–369.

He, X., Koo, B.B., Killiany, R.J., 2016. Edited Magnetic Resonance Spectroscopy Detects an Age-Related Decline in Nonhuman Primate Brain GABA Levels. Biomed Res Int 2016, 6523909.

Hillebrand, A., Barnes, G.R., Bosboom, J.L., Berendse, H.W., Stam, C.J., 2012. Frequency-dependent functional connectivity within resting-state networks: an atlas-based MEG beamformer solution. NeuroImage 59, 3909–3921.

Hua, T., Kao, C., Sun, Q., Li, X., Zhou, Y., 2008. Decreased proportion of GABA neurons accompanies age-related degradation of neuronal function in cat striate cortex. Brain Res Bull 75, 119–125.

Kuhlmann, L., Foster, B.L., Liley, D.T., 2013. Modulation of functional EEG networks by the NMDA antagonist nitrous oxide. PLoS One 8, e56434.

Lai, M.C., Lombardo, M.V., Chakrabarti, B., Sadek, S.A., Pasco, G., Wheelwright, S.J., Bullmore, E.T., Baron-Cohen, S., Consortium, M.A., Suckling, J., 2010. A shift to randomness of brain oscillations in people with autism. Biol Psychiatry 68, 1092–1099.

Leventhal, A.G., Wang, Y., Pu, M., Zhou, Y., Ma, Y., 2003. GABA and its agonists improved visual cortical function in senescent monkeys. Science 300, 812–815.

Liley, D.T., Cadusch, P.J., Dafilis, M.P., 2002. A spatially continuous mean field theory of electrocortical activity. Network 13, 67–113.

Liley, D.T., Sinclair, N.C., Lipping, T., Heyse, B., Vereecke, H.E., Struys, M.M., 2010. Propofol and remifentanil differentially modulate frontal electroencephalographic activity. Anesthesiology 113, 292–304.

Linden, H., Pettersen, K.H., Einevoll, G.T., 2010. Intrinsic dendritic filtering gives low-pass power spectra of local field potentials. J Comput Neurosci 29, 423–444.

Maxim, V., Sendur, L., Fadili, J., Suckling, J., Gould, R., Howard, R., Bullmore, E., 2005. Fractional Gaussian noise, functional MRI and Alzheimer's disease. Neuroimage 25, 141–158.

Miller, K.J., Sorensen, L.B., Ojemann, J.G., den Nijs, M., 2009. Power-law scaling in the brain surface electric potential. PLoS Comput Biol 5, e1000609.

Milotti, E., 2002. 1/f noise: a pedagogical review. arXiv preprint physics/0204033.

Muthukumaraswamy, S.D., 2013. High-frequency brain activity and muscle artifacts in MEG/EEG: a review and recommendations. Frontiers in Human Neuroscience 7, 138.

Muthukumaraswamy, S.D., Routley, B., Droog, W., Singh, K.D., Hamandi, K., 2016. The effects of AMPA blockade on the spectral profile of human early visual cortex recordings studied with non-invasive MEG. Cortex 81, 266–275.

Muthukumaraswamy, S.D., Shaw, A.D., Jackson, L.E., Hall, J.E., Singh, K.D., Saxena, N., 2015. Evidence that subanaesthetic doses of ketamine cause sustained disruptions of NMDA and AMPA-mediated frontoparietal connectivity in humans. Journal of Neuroscience 35, 11694–11706.

Nagasaka, Y., Shimoda, K., Fujii, N., 2011. Multidimensional recording (MDR) and data sharing: an ecological open research and educational platform for neuroscience. PLoS One 6, e22561.

Nolte, G., 2003. The magnetic lead field theorem in the quasi-static approximation and its use for magnetoencephalography forward calculation in realistic volume conductors. Physics in Medicine and Biology 48, 3637–3652.

Nutt, D., Wilson, S., Lingford-Hughes, A., Myers, J., Papadopoulos, A., Muthukumaraswamy, S., 2015. Differences between magnetoencephalographic (MEG) spectral profiles of drugs acting on GABA at synaptic and extrasynaptic sites: A study in healthy volunteers. Neuropharmacology 88, 155–163.

Oostenveld, R., Fries, P., Maris, E., Schoffelen, J.M., 2011. FieldTrip: Open source software for advanced analysis of MEG, EEG, and invasive electrophysiological data. Comput Intell Neurosci 2011, 156869.

Perrin, F., Pernier, J., Bertrand, O., Echallier, J.F., 1989. Spherical splines for scalp potential and current density mapping. Electroencephalography and Clinical Neurophysiology 72, 184–187.

Peterson, E.J., ROsen, B.Q., Campbell, A.M., Belger, A., Voytek, B., 2017. 1/f neural noise is a better predictor of schizophrenia than neural oscillations. BioRxiV.

Pettersen, K.H., Einevoll, G.T., 2008. Amplitude variability and extracellular low-pass filtering of neuronal spikes. Biophys J 94, 784–802.

Porges, E.C., Woods, A.J., Edden, R.A., Puts, N.A., Harris, A.D., Chen, H., Garcia, A.M., Seider, T.R., Lamb, D.G., Williamson, J.B., Cohen, R.A., 2017. Frontal Gamma-Aminobutyric Acid Concentrations Are Associated With Cognitive Performance in Older Adults. Biol Psychiatry Cogn Neurosci Neuroimaging 2, 38–44.

Pritchard, W.S., 1992. The brain in fractal time: 1/f-like power spectrum scaling of the human electroencephalogram. Int J Neurosci 66, 119–129.

Robinson, P.A., Rennie, C.J., Rowe, D.L., 2002. Dynamics of large-scale brain activity in normal arousal states and epileptic seizures. Phys Rev E Stat Nonlin Soft Matter Phys 65, 041924.

Schreiber, T., Schmitz, A., 1996. Improved Surrogate Data for Nonlinearity Tests. Phys Rev Lett 77, 635–638.

Sleigh, J., Harvey, M., Voss, L., Denny, B., 2014. Ketamine – More mechanisms of action than just NMDA blockade. Trends in Anaesthesia and Critical Care 4, 76–81.

Tzourio-Mazoyer, N., Landeau, B., Papathanassiou, D., Crivello, F., Etard, O., Delcroix, N., Mazoyer, B., Joliot, M., 2002. Automated anatomical labeling of activations in SPM using a macroscopic anatomical parcellation of the MNI MRI single-subject brain. NeuroImage 15, 273–289.

Van Veen, B.D., van Drongelen, W., Yuchtman, M., Suzuki, A., 1997. Localization of Brain Electrical Activity via Linearly Constrained Minimum Variance Spatial Filtering. Ieee Transactions on Biomedical Engineering 44, 867–880.

Voytek, B., Kramer, M.A., Case, J., Lepage, K.Q., Tempesta, Z.R., Knight, R.T., Gazzaley, A., 2015. Age-Related Changes in 1/f Neural Electrophysiological Noise. J Neurosci 35, 13257–13265.

Vrba, J., Robinson, S.E., 2001. Signal processing in magnetoencephalography. Methods 25, 249–271.

Walter, G., 1957. The Brain as a Machine. Proceedings of the Royal Society of Medicine 50, 799–808.

Wei, M., Qin, J., Yan, R., Li, H., Yao, Z., Lu, Q., 2013. Identifying major depressive disorder using Hurst exponent of resting-state brain networks. Psychiatry Research: Neuroimaging 214, 306–312.

Wen, H., Liu, Z., 2016a. Broadband Electrophysiological Dynamics Contribute to Global Resting-State fMRI Signal. J Neurosci 36, 6030–6040.

Wen, H., Liu, Z., 2016b. Separating Fractal and Oscillatory Components in the Power Spectrum of Neurophysiological Signal. Brain Topogr 29, 13–26.

Wink, A.M., Bernard, F., Salvador, R., Bullmore, E., Suckling, J., 2006. Age and cholinergic effects on hemodynamics and functional coherence of human hippocampus. Neurobiol Aging 27, 1395–1404.

Yamamoto Y, RL, H., 1993. Extracting fractal components from time series. Physica D-Nonlinear Phenomena 68, 250–264.

Yanagawa, T., Chao, Z.C., Hasegawa, N., Fujii, N., 2013. Large-scale information flow in conscious and unconscious states: an ECoG study in monkeys. PLoS One 8, e80845.

